# Genome-edited zebrafish model of *ABCC8* loss-of-function disease

**DOI:** 10.1101/2021.10.10.461291

**Authors:** Jennifer M. Ikle, Robert C. Tryon, Soma S. Singareddy, Nathaniel W. York, Maria S. Remedi, Colin G. Nichols

**Affiliations:** Department of Cell Biology and Physiology, Washington University in St. Louis, St. Louis, Missouri, United States; Center for the Investigation of Membrane Excitability Diseases, Washington University in St. Louis School of Medicine, St. Louis, Missouri, United States; Department of Pediatrics, Washington University in St. Louis School of Medicine, St. Louis, Missouri, United States; Department of Medicine, Division of Endocrinology, Metabolism, and Lipid Research, Washington University in St. Louis School of Medicine, St. Louis, Missouri, United States

**Keywords:** K_ATP_, Metabolism, Pancreas, Zebrafish, Calcium Channels, Insulin Secretion

## Abstract

K_ATP_ channel gain- (GOF) and loss-of-function (LOF) mutations in SUR1 or Kir6.2 underlie human neonatal diabetes mellitus (NDM) and congenital hyperinsulinism (CHI), respectively. Genetically modified mice with transgenic overexpression of GOF mutations recapitulate many features of human NDM and transgenic mice expressing incomplete K_ATP_ LOF do reiterate mild hyperinsulinism, although complete K_ATP_ knockout mice do not exhibit persistent hyperinsulinism. We have shown that islet excitability and glucose homeostasis in zebrafish are regulated by identical K_ATP_ channels to those in mammals, and to explore the possibility of using zebrafish for modeling CHI, we have examined SUR1 truncation mutation (K499X) introduced into *the abcc8* gene. Patch-clamp analysis confirmed complete absence of channel activity in β-cells from K499X (SUR1^-/-^) fish. No difference in random blood glucose was detected in heterozygous SUR1^+/-^ fish, nor in homozygous SUR1^-/-^ fish, mimicking findings in SUR1 knockout mice. Mutant fish also demonstrated impaired glucose tolerance, similar to LOF mouse models. In paralleling features of mammalian hyperinsulinism resulting from loss-of-function mutations, this gene-edited animal provides a valid zebrafish model of K_ATP_ LOF driven-dependent pancreatic disease.

**Key Points:**

1. Loss-of function in the Kir6.2 (KCNJ11) and SUR1 (ABCC8)-encoded pancreatic islet β-cell K_ATP_ channels underlie congenital hyperinsulinism. Mouse models reiterate key features, but zebrafish models could provide a powerful model for further analysis and therapy testing.
2. An early nonsense mutation in exon 10 of SUR1 was generated by ENU mutagenesis.
3. Patch-clamp analysis revealed an absence of β-cells of SUR1 truncation mutants.
4. Ca imaging demonstrated elevated basal [Ca]_i_ in β-cells with SUR1 truncation.
5. Homozygous SUR1 truncation mutants had normal fasting glucose but impaired glucose tolerance as adults, mimicking findings in mouse SUR1 knockouts.
6. In paralleling features of mammalian diabetes and hyperinsulinism resulting from equivalent loss-of-function mutations, this gene-edited animal provides a valid zebrafish model of K_ATP_ -LOF dependent pancreatic diseases.

## Introduction

Electrical activity is a key component of insulin secretion from β-cells^1^ and is critically regulated by ATP-sensitive potassium (K_ATP_) channels. In mammals, pancreatic K_ATP_ channels are composed of four SUR1 subunits (encoded by *ABCC8*) and four Kir6.2 subunits (encoded by *KCNJ11*) ^2,3^. At low plasma [glucose], K_ATP_ channels are normally open, the cell membrane is hyperpolarized, and voltage-dependent calcium channels (VDCCs) are closed, thus inhibiting insulin secretion ^4^. Glucose metabolism increases the intracellular [ATP]/[ADP] ratio via enhanced β-cell glycolysis and oxidative phosphorylation. This causes closure of the K_ATP_ channels, leading to membrane depolarization, calcium influx through VDCCs, and triggering of insulin release ^5,6^. Congenital hyperinsulinism (CHI) is the most common cause of hypoglycemia in neonates and infants^7^ and is often linked to loss-of-function (LOF) mutations in either *KCNJ11* or *ABCC8* ^*8*^, which result in reduced K_ATP_ channel activity, β-cell hyperexcitability, and excessive insulin secretion ^9^. Mice with transgenic expression of K_ATP_ LOF mutations, as well as mice with heterozygous *KCNJ11* or *ABCC8* gene knockout reiterate persistent hyperinsulinism ^15,16^. However, homozygous K_ATP_ knockout mice do not exhibit persistent hyperinsulinism, instead, they exhibit an unexplained loss of insulin secretion and glucose intolerance ^17-20^.

Therapeutic approaches to, and management of, CHI will benefit from novel animal models and new insights into disease processes, leading the way to new opportunites for treatment. We have shown that K_ATP_ channels are expressed in β-cells within the zebrafish (*Danio rerio*) islet, that they are functionally similar to their mammalian orthologues ^21^, and that they exert similar glucose-dependent control of intracellular [Ca] ([Ca]_i_) ^22,23^. Activation of these channels by the drug diazoxide^21^ or overexpression of ATP-insensitive transgenes in β-cells ^23^ can similarly alter metabolic response to glucose. To provide a model of K_ATP_ LOF, we have investigated a zebrafish model of loss-of-function of K_ATP_ in which an early nonsense mutation, predicted to lead to premature truncation of SUR1, was introduced into *abcc8*. This zebrafish mutant recapitulates key features of human K_ATP_ LOF, and provides a model for further analysis and testing of potential therapeutics, that may facilitate advances in clinical management and help identify new therapies, by providing a high throughput platform for understanding mechanisms and testing potential therapeutic approaches.

## Results

### Genome-modified zebrafish model of SUR1 LOF

A number of K_ATP_ mutations have been described to cause CHI, with the causal mutation being in the SUR1 subunit more often than in Kir6.2^9,24^. Although gating mutations are a common underlying cause, mutations that result in loss of functional protein or failure to traffic to the cell membrane, are also prominent^25,26^. To model loss of functional protein, we developed an *abcc8* mutant line that was originally generated by ENU mutagenesis, from the Zebrafish Mutation Project^27^. These fish contain an early stop codon (X) mutation in exon 10 of SUR1 (K499X), which results in disruption of transmembrane domain 1 (Fig. 1A), and is predicted to result in loss of functional SUR1 protein. Complete knockout of channel activity was confirmed in isolated membrane patches of β-cells from homozygous mutant fish (see below), which we thus term SUR1^-/-^. Infants with CHI often have macrosomia^28,29^, but, as with mouse knockouts^18,30,31^ there was no significant difference in growth between SUR1 mutants and wild type (Fig. 1B). In mice with such a marked loss of K_ATP_ channels, a common theme of early transient hyperinsulinemic hypoglycemia followed by normoglycemia or impaired glucose tolerance as adults has been described ^16,18,19,31^. A similar progression has also been seen in some humans with CHI due to K_ATP_ LOF ^25,32,33^. Preliminary measurements suggest that SUR1^-/-^ larvae also have lower whole-body glucose compared to wildtype (not shown), but blood glucose in both homo- and heterozygous fish was not different from WT (Fig. 1C).

**Figure 1.**
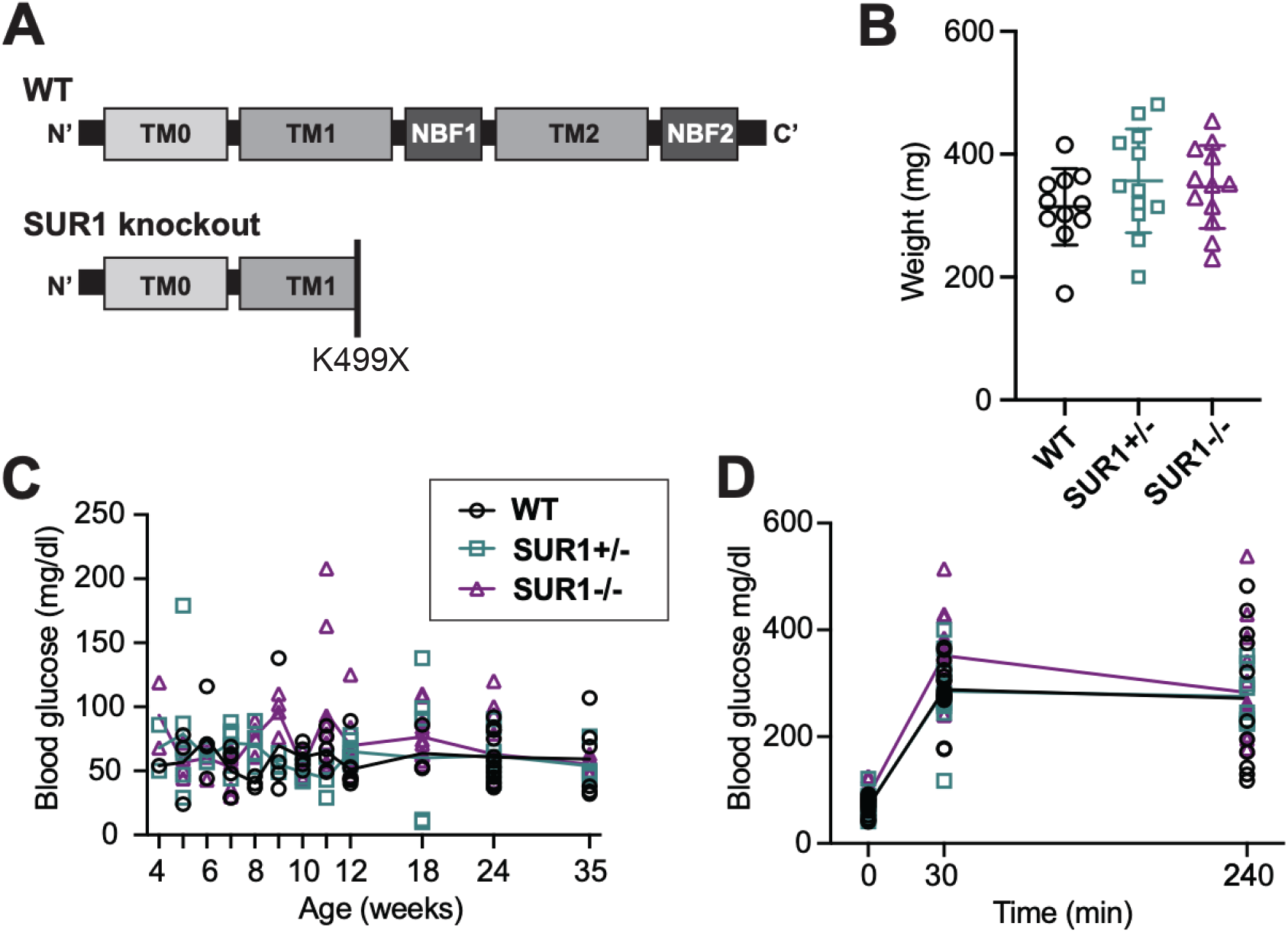
Genome-modified zebrafish model of SUR1 LOF. (A) Schematic of SUR1 protein structure showing functional domains and predicted truncation resulting from premature stop mutation (K4899X) within TMD1. (B) Body weight in WT (n=11), heterozygous SUR1^+/-^ (n=12) and SUR1^-/-^ (n=12) fish at age 8 weeks. (C) Fasting blood glucose over time shows no significant difference between homozygous mutants and controls (n=11-16 per timepoint). (D) Intraperitoneal GTT shows non-significantly impaired response to glucose load in SUR1^-/-^, relative to WT or SUR1^+/-^ fish (n=8-16 per timepoint).

Intraperitoneal glucose tolerance test (IPGTT) was performed on adult zebrafish. We have previously shown that IPGTT in wild-type zebrafish have a peak in blood glucose around 30 minutes with a slow return to normal over four to six hours ^21^. Here, we found higher peak glucose in SUR1^-/-^ compared to wild type, but similar return to baseline by 4 hours post injection (Fig. 1D). This parallels the findings in mouse SUR1 and Kir6.2 knockout models of complete loss of K_ATP_, which lack persistent hypoglycemia and instead exhibit glucose intolerance and loss of insulin secretion as adults ^17-20^.

### Molecular consequences of introduced gene modifications

We crossed SUR1 (K499X) fish to *ins:GFP* expressing fish to assess morphology and insulin gene promoter activity. GFP fluorescence was used to identify β-cells in isolated islets via confocal microscopy. There was no obvious deficit of β -cell density or GFP density in SUR1 mutants compared to WT (not shown). Inside-out patch clamp recordings from β-cells isolated from primary islets of SUR1^-/-^ and wild-type fish, both containing *ins:GFP* transgene, were also identified by the presence of green fluorescence. Excised inside-out patch-clamp experiments (Fig. 2A,B) confirmed effective knockout, with complete absence of K_ATP_ channels in SUR1 homozygous mutant β-cells. These recordings additionally confirmed no responsivity to the channel opener diazoxide, consistent with a complete loss of SUR1-dependent K_ATP_ channel function.

**Figure 2.**
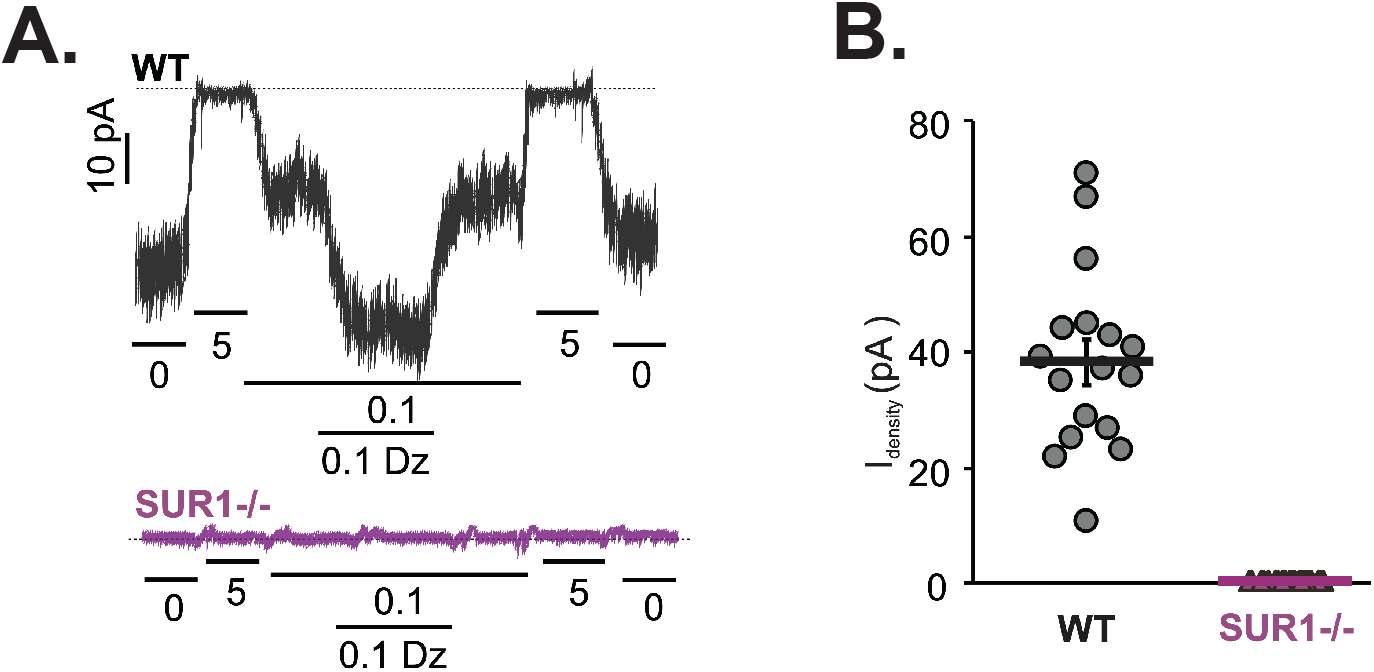
SUR1 [499X] abolishes K_ATP_ channel activity. (A) Representative K_ATP_ channel activity in inside-out patch clamp recordings from inside-out patch clamp recordings of WT (above) and homozygous SUR1[499X] (SUR1^-/-^) mutant β-cells (below). Voltage was clamped at -50mV. (B) K_ATP_ channel density in WT and SUR1-/-patches (n=17,10).

### Excitability consequences of introduced gene modifications

K_ATP_ channel LOF is predicted to increase islet excitability and increase Ca entry into β-cells. SUR1^-/-^ fish were crossed with transgenic GCaMP6s fish carrying *Tg(ins:GCaMP6s)* ^34^, and time-lapse confocal fluorescent microscopy was carried out on *ex vivo* perifused whole adult islets. Consistent with prior findings, controls showed significant increase in relative fluorescence when transitioned from low (2 mM) to high (20 mM) glucose (Fig. 3A, B). In SUR1^-/-^ zebrafish, carrying the same *Tg(ins:GCaMP6s)*, [Ca] imaging revealed elevated basal fluorescence in low (2 mM) glucose concentration, reflective of basal depolarization in mutant β-cells, but no significant further increase in high (20 mM) glucose (Fig. 3A,B). These results are consistent with loss of K_ATP_ in SUR1^-/-^ causing electrical activity at low [glucose] by depolarizing cells, and a failure to respond appropriately to subsequent increases in [glucose].

**Figure 3.**
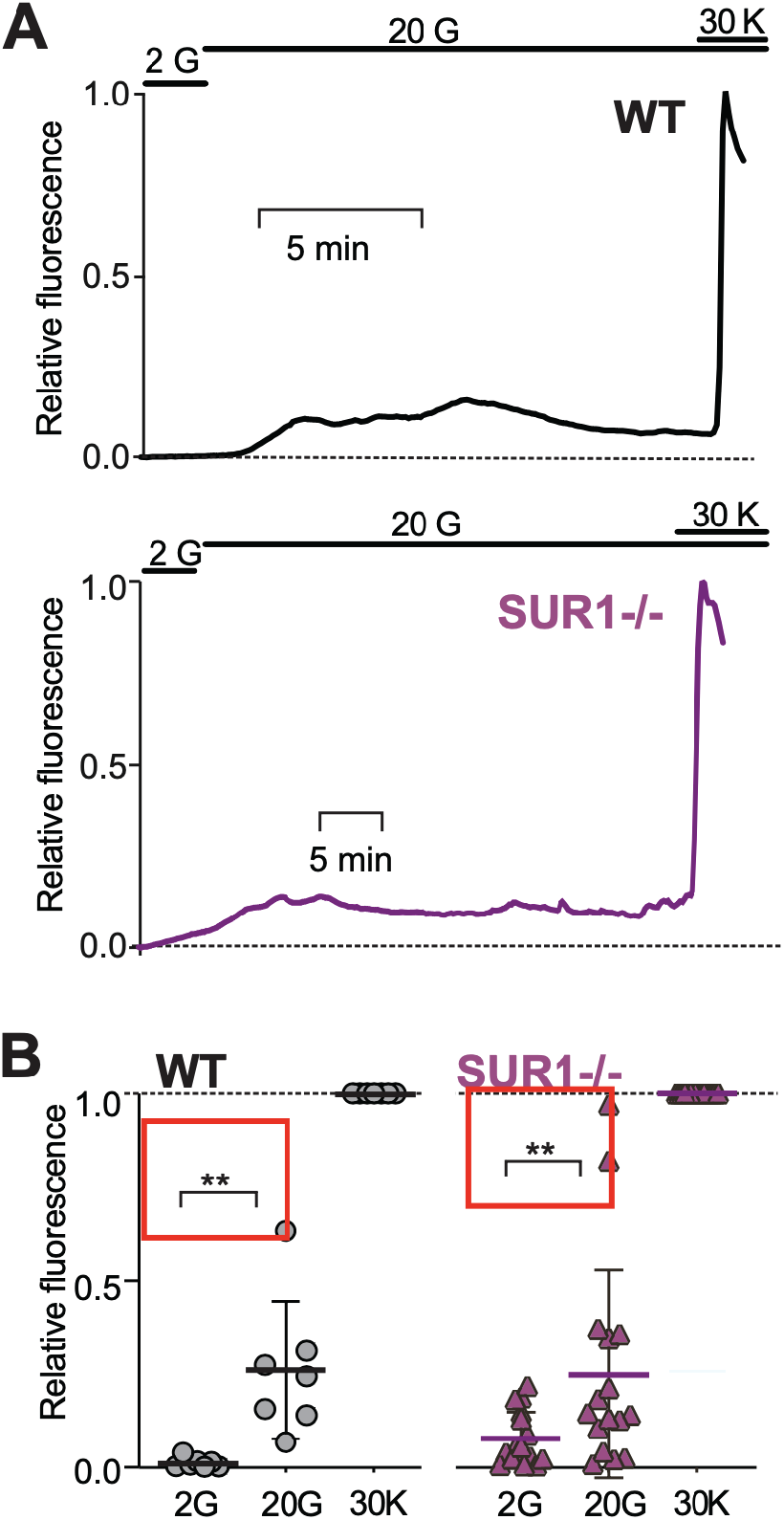
SUR1^-/-^ islets exhibit elevated basal [Ca] and reduced responsivity to glucose. (A) Representative recordings of intracellular calcium in the presence of 2 mM glucose (2G) and following switch to 20 mM glucose (20G) and then 30 mM KCl (30K), for WT (*above*) and SUR1^-/-^ (*below*). Fluorescence is normalized to maximum fluorescence in 30K (*f* = 1), and minimum fluorescence (*f*=0) anywhere within the record. (B) Average fluorescence in each condition for WT (N=8) and SUR1^-/-^ (n= 14). Data in B are analyzed by 1-way ANOVA followed by multiple unpaired t-tests. (*) p <0.05, (**) p <0.01.

## Discussion

It is likely that many human HI mutations will cause only incomplete loss of K_ATP_ channel activity ^35,36^, and both active K_ATP_ channels and sensitivity to the K_ATP_ channel opener diazoxide have been detected in some K_ATP_ -dependent HI patients ^37,38^. Mice with *partial* loss of K_ATP_ activity mimic this hyperinsulinemic phenotype, and secrete insulin at lower [glucose] than controls ^16^. However, while mice with *complete* loss of K_ATP_ exhibit elevated serum insulin and hypoglycemia in the neonatal period ^16,18,19,31^, they then rapidly develop hyperglycemia with reduced insulin secretion, a phenomenon that persists through adulthood. This cross-over to loss of secretion, in the face of continual excitation, reflects a marked down-regulation of the secretory process itself. The underlying cause remains elusive, and thus far it is unknown whether this is a mouse-specific progression, or reflects processes that may also be involved in human HI. Here, we demonstrate a similar glucose-intolerant and non-hypoglycemic phenotype in zebrafish that completely lack K_ATP_ channels. The premature stop mutation at position 499 is expected to result in a severely truncated protein, containing only the TMD0 region and half of the second TMD1 region, and lacking both nucleotide binding folds (NBFs). Previous studies have shown that essentially no functional channels are formed when NBF1 is absent ^39^ and, accordingly, no channels were identified in patches from isolated these SUR1^-/-^ β-cells. Consistent with findings in SUR1^-/-^ mice, blood glucose was essentially normal in adult SUR1^-/-^ fish, but glucose tolerance was reduced. These fish thus confirm a common finding from fish to mouse and may provide a useful model for further exploring the unexplained phenomenon of glucose intolerance and even progression to diabetes that is seen in K_ATP_ –LOF mice and K_ATP_-dependent CHI patients.

## Conclusions

In paralleling progressive crossover to glucose intolerance that is seen in congenital hyperinsulinism resulting from loss-of-function mutations, these gene-edited animals provide a valid zebrafish model of K_ATP_ LOF dependent pancreatic diseases.

## Abbreviations

KCNJ11: potassium voltage-gated channel subfamily J member 11
ABCC8: ATP Binding Cassette Subfamily C Member 8
Kir6.2: ATP-sensitive potassium channel pore subunit, inward-rectifying
SUR1: Sulfonylurea receptor 1
VDCC: Voltage-dependent calcium channel
NDM: Neonatal diabetes mellitus 1

## Author contributions

CGN and JMI conceived the study; JMI, RCT, SSS, NWY, LY carried out the experiments; JMI, RCT, SSS, NWY analyzed the data; CGN, MSR, JMI, RCT, SSS, NWY interpreted the results; MSR and JMI participated in the design of experiments. JMI and CGN wrote the paper, which was edited by other authors. All authors gave final approval for the manuscript.

## Acknowledgements and Funding

We would like to acknowledge the assistance of the Washington University in St Louis Zebrafish Facility (http://zebrafishfacility.wustl.edu/).

This work was supported by grants from the National Institutes of Health to CGN (R01 DK109407), MSR (R01 DK 098584). JMI was supported by NIH T32HD043010-15, Pediatric Endocrine Society, and Endocrine Fellows Foundation. This work was additionally supported by the Washington University in St. Louis Diabetes Research Center (DK 020579) and the Washington University Center for Cellular Imaging (ZRSC 305636608M to MSR).

## Ethics statement

All procedures were approved by the Washington University Institutional Animal Care and Use Committee.

## Competing interests

The authors declare no competing interests.

## Methods

### Ethical Approval

All animal procedures were approved by the Washington University in St. Louis Institutional Animal Care and Use Committee.

### SUR1 ENU generated nonsense mutation

ENU-mutagenesis was performed at the Sanger Institute, as part of the Zebrafish Mutation Project, using N-ethyl-N-nitrosourea (ENU) mutagenesis to attempt to identify knockout alleles for all protein-coding regions in the zebrafish genome (https://www.sanger.ac.uk/resources/zebrafish/zmp/). This project outcrosses ENU-mutagenized F0 males to create a population of F1 fish heterozygous for ENU-induced mutations, which were then obtained through the Zebrafish International Research Consortium (ZIRC). The *abcc8(sa15863)* nonsense mutant allele (K499-STOP, TTCTGGCTCCRGTGCAGTACTTTGTGGCAACCAAGTTATCAGATGCACAG**[A>T]**AAAG CACATTGGTGAGCTACTTTATTTTGGTTAATGTCCTAATGAGGCCA) was obtained from the Zebrafish Mutation Project^27^, through Zebrafish International Resource Center. Homozygous K499-STOP mutants were generated by in-crossing heterozygous carriers and the progeny were genotyped by Transnetyx using restriction digest with the inserted digestion site for HpyCHRIII which is inserted into the mutant allele (Forward primer: TTGTTGTTGTCTGCTTTTTGC. Reverse primer: TTTACAAGCACAGCGCTCAC) to identify homozygotes.

### Animal lines and maintenance

In addition to the mutant lines above, we used AB wild-type fish as well as previously described β-cell-specific GCaMP6s-expressing transgenic fish *Tg(−1*.*0ins:gCaMP6s)*^*stl44134*^ and *Tg(−1*.*0ins:eGFP)*^*sc1*^ ^40^. Wild-type controls were on the AB background. All fish lines were housed in the Washington University Zebrafish Facility under standard conditions, the details of which can be found at: http://zebrafishfacility.wustl.edu/documents.html.

### Electrophysiological analyses

Islets were isolated from zebrafish, and single β-cells were dissociated, as previously described and recordings were performed on GFP-positive cells^21^. Excised patch recordings were performed using pipettes with a resistance of 1-2 MΩ when filled with pipette solution. Bath and pipette solution (K-INT) contained (in mM): 140 KCl, 10 HEPES, 1 EGTA (pH 7.4 with KOH). All recordings were performed at -50mV holding potential and absence or presence of nucleotides was adjusted in bath solution as indicated. Data were tested for statistical significance using Welch’s *t*-test. A *p* value of <0.05 was considered significant.

### Blood glucose measurements and glucose tolerance test

Zebrafish were fasted for 18-20 hours prior to blood glucose measurement. Blood glucose was measured as described^21^. Intraperitoneal glucose tolerance test was performed as described^21^ on similarly fasted zebrafish.

### Growth measurements

To obtain growth data, fish were briefly anesthetized in tricaine, excess water removed with Kimwipe, then weighed on a digital scale with precision to 1 μg, before returning fish to fresh water to recover from anesthesia.

### Chemicals

All salts, amino acids, and other compounds were purchased from Sigma, except where indicated above.

### Statistics

Statistical analyses were performed in GraphPad prism. For multiple group column data comparisons, data were analyzed by unpaired t-tests where normality assumptions were met. All values are indicated as mean ± SEM, except where noted. For dose-response curves, data were fitted using nonlinear regression.

## Notes

### Competing Interest Statement

The authors have declared no competing interest.

